# Prenatal Diagnosis of Fetuses with Increased Nuchal Translucency by Genome Sequencing Analysis

**DOI:** 10.1101/667311

**Authors:** Kwong Wai Choy, Huilin Wang, Mengmeng Shi, Jingsi Chen, Zhenjun Yang, Rui Zhang, Huanchen Yan, Yanfang Wang, Shaoyun Chen, Matthew Hoi Kin Chau, Ye Cao, Olivia YM Chan, Yvonne K Kwok, Yuanfang Zhu, Min Chen, Tak Yeung Leung, Zirui Dong

## Abstract

**Background:** Increased Nuchal Translucency (NT) is an important biomarker associated with increased risk of fetal structural anomalies. It is known to be contributed by a wide range of genetic etiologies from single nucleotide variants to those affecting millions of base-pairs. Currently, prenatal diagnosis is routinely performed by karyotyping and chromosomal microarray analysis (CMA), however, both of them have limited resolution. The diversity of the genetic etiologies warrants an integrated assay such as genome sequencing (GS) for comprehensive detection of genomic variants. Herein, we aim to evaluate the feasibility of applying GS in prenatal diagnosis for the fetuses with increased NT.

**Methods:** We retrospectively applied GS (>30-fold) for fetuses with increased NT (≥3.5-mm), who underwent routine prenatal diagnosis. Detection of single-nucleotide variants, copy-number variants and structural rearrangements was performed simultaneously and the results were integrated for interpretation in accordance with the guidelines of the American College of Medical Genetics and Genomics. Pathogenic or likely pathogenic (P/LP) variants were selected for validation and parental confirmation, when available.

**Results:** Overall, 50 fetuses were enrolled, including 34 cases with isolated increased NT and 16 cases with other fetal structural malformations. Routine CMA and karyotyping reported eight P/LP CNVs, yielding a diagnostic rate of 16.0% (8/50). In comparison, GS provided a 2-fold increase in diagnostic yield (32.0%, 16/50), including one mosaic turner syndrome, eight cases with microdeletions/microduplications and seven cases with P/LP point mutations. Moreover, GS identified two cryptic insertions and two inversions. Follow-up study further demonstrated the potential pathogenicity of an apparently balanced insertion which disrupted an OMIM autosomal dominant disease-causing gene at the inserted site.

**Conclusions:** Our study demonstrates that applying GS in fetuses with increased NT can comprehensively detect and delineate the various genomic variants that are causative to the diseases. Importantly, prenatal diagnosis by GS doubled the diagnostic yield compared with routine protocols. Given a comparable turn-around-time and less DNA required, our study provides strong evidence to facilitate GS in prenatal diagnosis, particularly in fetuses with increased NT.

## Introduction

Detection of fetuses with increased nuchal translucency (NT) in routine first-trimester ultrasound screening has been widely used as a sensitive indication for fetal chromosomal abnormalities and/or fetal structural anomalies, such as congenital heart disorders or neurodevelopmental anomalies detected in later gestations (Leung et al., 2011;Huang et al., 2014;Socolov et al., 2017;Sinajon et al., 2019). Fetuses with increased NT and structural malformations are frequently contributed by genetic abnormalities and have poor prognoses. However, more than 80% of such cases do not obtain a causative result with the current routine prenatal diagnostic tests (Leung et al., 2011;Huang et al., 2014;Yang et al., 2017;Sinajon et al., 2019), challenging genetic counseling and clinical management. In addition, pathogenic copy-number variants (CNVs) only account for 0.8 to 5.3% of these fetuses with isolated increased NT (with/without other soft markers) (Leung et al., 2011;Huang et al., 2014), and part of these cases would also have poor outcomes. Therefore, a test for the comprehensive detection of disease associated genomic variants including numerical disorders, structural rearrangements, CNVs and point mutations in this prenatal cohort is warranted.

In prenatal diagnosis, quantitative fluorescent PCR (QF-PCR) is routinely conducted for the detection of maternal cell contamination (MCC) and common aneuploidy [such as Trisomy 21 (Choy et al., 2014;Sinajon et al., 2019)]. In addition, since 2010, chromosomal microarray analysis (CMA) has been recommended as the first-tier test for high-risk pregnancies in identification of microscopic or submicroscopic CNVs (Leung et al., 2011;Huang et al., 2014;Sinajon et al., 2019). However, this approach is limited by its resolution and it cannot detect single nucleotide variants (SNVs) and small insertions/deletions (InDels). Owing to the breakthrough of molecular technologies such as next-generation sequencing and its reduction of costs over the years, whole-exome sequencing (or exome sequencing, WES) has been applied for both research purposes and clinical use (Drury et al., 2015;Fu et al., 2018;Leung et al., 2018;Normand et al., 2018;Lord et al., 2019;Petrovski et al., 2019). Emerging studies show WES has the ability to provide genetic diagnoses ranging from 9.1% to 32% for the fetuses with a structural anomaly (Drury et al., 2015;Fu et al., 2018;Leung et al., 2018;Normand et al., 2018;Lord et al., 2019;Petrovski et al., 2019), while among these cases, WES yielded diagnoses in 3.2% to 21% of the fetuses with increased NT with/without structural malformations (Drury et al., 2015;Lord et al., 2019;Petrovski et al., 2019). However, most of these studies were conducted on prenatal cohorts after the exclusion of abnormal karyotypes and/or CMA results attributed to the cost and the limited ability of WES in CNV detection (Belkadi et al., 2015). These studies show the clinical utility of WES and CMA in prenatal diagnosis and warrants a combination of these two approaches for each case. Meanwhile, both WES and CMA are unable to detect apparently balanced structural rearrangements (or structural variants, SVs), a common limitation of the current methods but some of these rearrangements have been demonstrated to be disease-causing (Talkowski et al., 2012a). The wide spectrum of genetic etiologies in fetuses with increased NT ranging from single-base mutations to those affecting millions of base-pairs and numerical disorders, warrants a holistic approach for comprehensive detection of the disease-causing genetic variants.

Our previous studies have demonstrated the feasibility and potential diagnostic utility of applying low-pass whole-genome sequencing (or genome sequencing, GS) analysis in the detection of CNVs (Dong et al., 2016;Dong et al., 2017) and chromosomal structural rearrangements (Dong et al., 2014) including balanced translocations and inversions in both clinical cohorts and presumably normal populations in the 1000 Genomes Project (Dong et al., 2018a;Dong et al., 2018b). By increasing the read-depth to a minimal of 30-fold for the purpose of including SNV/InDel detection, GS is able to provide comprehensive detection of various genomic variants, thus, providing a unique platform for gene discovery and potential clinical application. However, evaluation of its clinical utility is warranted.

Herein, we aimed to apply GS for the investigation of genetic contributions to fetuses with increased NT with/without structural malformations and to evaluate the possibility for its potential clinical application.

## Materials and Methods

### Ethics, consent and permissions

The study protocol was approved by the Ethics Committee of the Joint Chinese University of Hong Kong-New Territories East Cluster Clinical Research Ethics Committee (CREC Ref. No. 2016.713), Jinan University and Guangzhou Medical University. From year 2014 to 2018, 50 pregnant women, whose fetus was diagnosed with increased NT (≥ 3.5-mm) with/without structural malformations (Leung et al., 2011;Huang et al., 2014) and had undergone prenatal diagnosis by CMA (and karyotyping if available) after a negative finding from QF-PCR (Choy et al., 2014), were recruited in this study. The recruitment criteria included fetuses with increased NT detected and CMA results were available without the selection of (1) whether any fetal malformation was detected; (2) the results from CMA and/or karyotyping; (3) the timing for sample selection (CVS or AF) and (4) the pregnancy outcome (such as terminated pregnancies or livebirths) (Figure 1). Written informed consent was obtained from each participant for the purpose of this study and any findings from the genome sequencing would not be disclosed to the patients. Routine CMA results were available in all cases and 90.0% (45/50) of them also had G-banded chromosome analysis results. Among them, chorionic villus sampling (CVS) was conducted for 37 cases at the time of first-trimester ultrasound screening, while amniotic fluid (AF) was obtained for the other 13 cases during the second-trimester. Parental peripheral blood samples were collected at the time of prenatal sample retrieval if available.

**Figure 1.**
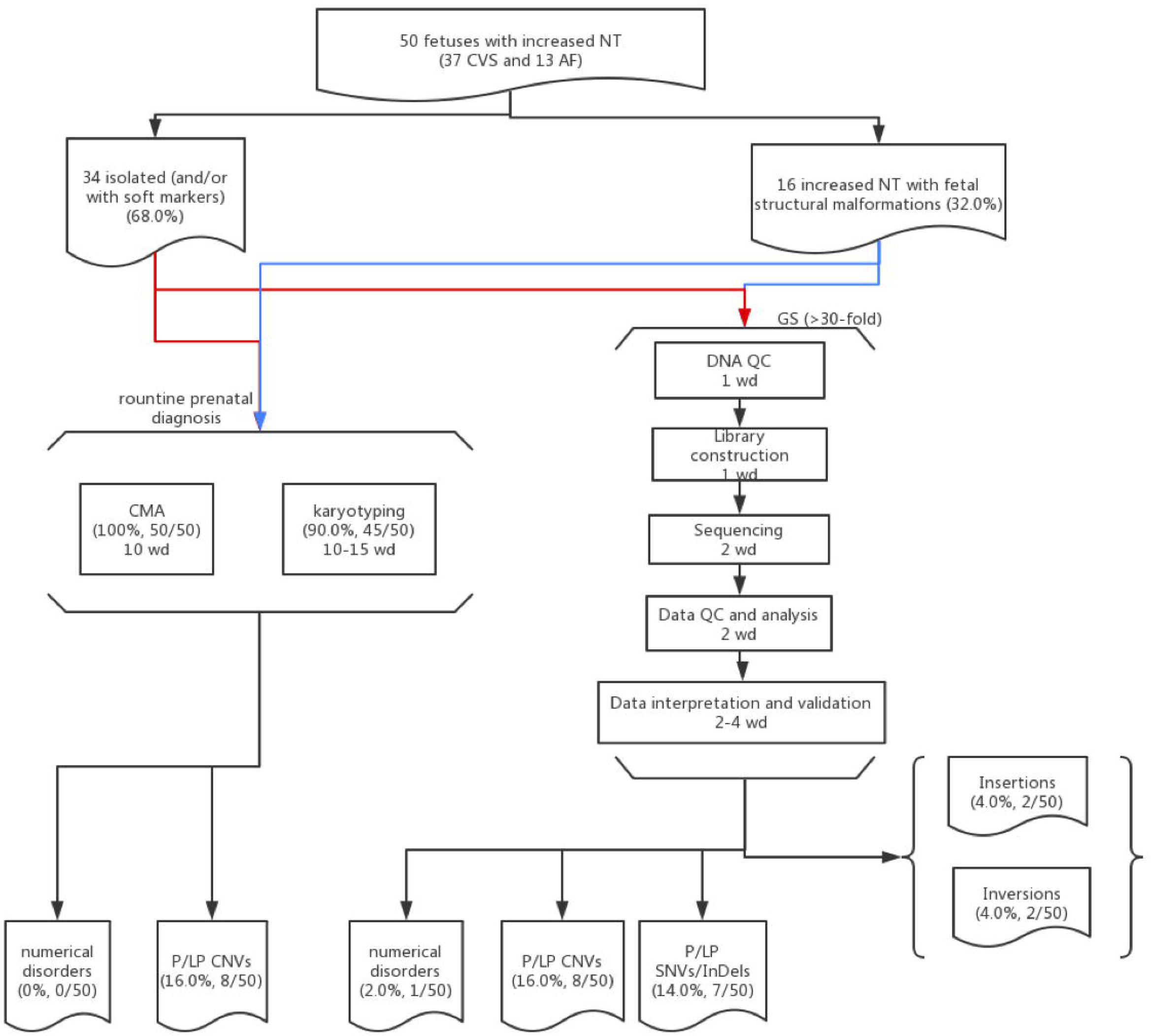
Flowchart of this Study. Detailed methods and results are described in the main text. 50 cases with isolated (red lines) or syndromic increased NT (blue lines) and prenatal diagnosis results available (CMA and karyotyping) were selected for GS. The detailed procedures with the estimated turn-around-time expressed as working day (wd) are provided in each box.

### DNA Preparation

Genomic DNA from chorionic villi or amniotic fluids (AF) were extracted using the DNeasy Blood & Tissue Kit (Cat No./ID: 69506, Qiagen, Hilden, Germany) at the time of routine CMA testing. DNA was quantified with the Quant-iT dsDNA HS Assay kit (Invitrogen, Carlsbad, CA), DNA integrity was assessed by agarose gel electrophoresis and subsequently subjected for QF-PCR.

### CMA and Karyotyping for Routine Prenatal Diagnosis

Two CMA platforms were routinely used in our three prenatal genetic diagnosis centers: CytoScan 750K (Applied Biosystems, Affymetrix, Inc., Santa Clara, CA) and a well- established 8×60K fetal DNA chip (Agilent Technologies, Santa Clara, CA).

For the CytoScan 750K SNP-based platform, 250-ng of DNA samples was required and the experiment was conducted based on the manufacturer’s protocol (Lin et al., 2015). Aberration analysis was performed with the ChAS 2.0 software (Affymetrix) (Lin et al., 2015). While for fetal DNA chip array-based comparative genomic hybridization (aCGH) platform (Leung et al., 2011), the experiment was conducted with a total of 300-ng DNA after treatment with RNase A (Qiagen) as input, and data analysis was performed by CytoGenomics according to the manufacturer’s protocol (Leung et al., 2011).

For 90.0% (45/50) cases, routine G-banded chromosome analysis of the CVS or AF was also conducted by certified the Medical Technologists/Cytogeneticists following the international guidelines [Standards and Guidelines for Clinical Genetics Laboratories (Section 5.1) of the American College of Medical Genetics and Genomics (ACMG) and Requirements for Cytogenetic Testing of the National Pathology Accreditation Advisory Council]. A minimum of two independent cultures were set up for each specimen and 15-20 metaphases were analyzed with two metaphases karyotyped at a resolution of 400 bands. In cases suspected of mosaicism, at least 30 metaphases were analyzed. Karyograms were interpreted by at least two Medical Technologists/Cytogeneticists and diagnoses were reported according to the International System for Human Cytogenomic Nomenclature (ISCN) 2016.

### GS for Fetal DNA

First, 100-ng of genomic DNA from each sample was sheared to fragment sizes ranging from 300∼500-kb by the Covaris S2 Focused Ultrasonicator (Covaris, Inc., Woburn, MA). Library construction including end-repairing, A-tailing and adapter-ligation and PCR amplification was conducted subsequently. The PCR products were then heat-denatured to form single strand DNAs, followed by circularization with DNA ligase. After construction of the DNA nanoballs, paired-end sequencing with 100-bp at each end was carried out for each sample with a minimal read-depth of 30-fold on the MGISEQ-2000 platform (BGI-Wuhan, Wuhan, China) (Huang et al., 2017).

### Data Analysis and Variants Detection

QC for the paired-end reads was assessed via FastQC (https://www.bioinformatics.babraham.ac.uk/projects/fastqc/) and subsequently aligned to the human reference genome (hg19) by Burrows-Wheeler Aligner (BWA) (Li and Durbin, 2009) and reformatted with SAMtools (Li et al., 2009). SNV and InDel detection was performed with HaplotypeCaller v3.4 from the Genome Analysis Toolkit (GATK, Broad Institute) (McKenna et al., 2010), and annotation by ANNOVAR (Wang et al., 2010) and InterVAR (Li and Wang, 2017) with public and our in-house databases.

CNV detection and SV analyses were performed by our previously published methods (Dong et al., 2014;Dong et al., 2016;Dong et al., 2018a) with uniquely aligned reads/read-pairs. For CNV analysis, all aligned reads were classified into each adjustable sliding window (50-kb with 5-kb increments) for identifying the candidate region(s) with CNVs, then they were classified again into each non-overlapping window (5-kb) for detection of the precise breakpoints with the module of Increment-Rate-of-Coverage. The rare CNVs (U-test P<0.0001) were then selected for further interpretation. For SV analysis, all chimeric read-pairs, which were aligned to different chromosomes or to the same chromosome but with a distance larger than expected (>10-kb), were selected for identification of translocations, inversions, insertions or complex rearrangements. Briefly, (1) Clustering: the chimeric read-pairs were clustered by sorting the aligned coordinates; (2) Systematic error filtering: each event was filtered against a control data set for the elimination of potential systematic errors; (3) Random error filtering: each event was filtered with a cluster property matrix with the reported parameters; and (4) Aligned orientations: each event was filtered based on p/q arm genetic exchange (joining type). Results of SNVs, CNVs and SVs analyses were integrated and reviewed for classification and interpretation of pathogenicity. The final results were also provided based on ISCN 2016.

### Data Interpretation and Validation

The clinical significance of the detected SNVs, InDels and CNVs were interpreted in accordance with the guidelines of the American College of Medical Genetics and Genomics (ACMG) and were classified into five categories: pathogenic, likely pathogenic, variant of uncertain significance, likely benign or benign. Prioritization of SNVs/InDels in each sample was based on the following criteria: (1) whether reported by ClinVAR or HGMD (human mutation gene database); (2) with a minor allele frequency ≤ 5% in the databases of ExAC (http://exac.broadinstitute.org) and gnomAD (https://gnomad.broadinstitute.org); (3) located in coding-region and exon-intron junctions; (4) with damaging/intolerant or splicing-change effect suggested by multiple biological algorithms (SIFT, Polyphen-2, MutationTaster, Human Splicing Finder and MaxEntScan); (5) located in an OMIM disease causing gene. For known mutations, correlation of ultrasound finding(s) with the reported phenotype(s) was conducted. For novel variants, the priority of further classification was conducted as (1) located gene was reported to be in autosomal dominant or X-linked dominant manner; (2) affecting gene was in autosomal recessive or X-lined recessive manner and a homozygous variant or more than one variant (suspected compound heterozygosity) were found. CNV interpretation was conducted based on our reported study (Dong et al., 2016). Potential disease-causing mutations were selected for validation and parental confirmation when available. No guidelines were available for SV interpretation, thus gene disruption or potential gene dysregulation by the disruption of regulatory elements or topological associated domains were used for further interpretation. Parental confirmation was conducted if available for determination of the mode of inheritance of the variants.

SNV/InDel/SV were validated by sanger sequencing. Genomic reference sequences (hg19) from each putative variant/breakpoint region were used for web-based primer design with Primer3 (http://primer3.ut.ee/) and NCBI Primer-Blast (http://www.ncbi.nlm.nih.gov/tools/primer-blast/). PCR amplification was performed with each pair of primers (**Supplementary Table S1**) in cases and control (in-house DNA from a presumably normal male subject) simultaneously. PCR products were sequenced by Sanger sequencing on an ABI 3730 machine (Applied Biosystems, Thermo Fisher Scientific, Wilmington, DE) and sequencing results were aligned with BLAT (https://genome.ucsc.edu/cgi-bin/hgBlat?command=start) for confirmation of the mutations or re-arranged DNA sequences (Dong et al., 2016).

For CNV validation, quantitative PCR (qPCR) was conducted for additional disease-causing CNVs identified by GS. Genomic reference sequences (hg19) of each deleted/duplicated region were used for web-based primer design with Primer3 and NCBI Primer-Blast. Melting curve analysis was carried out for each pair of primers, and the PCR efficiency ranging 95% to 105% was determined by using the standard curve method. Each reaction was performed in a 10-μl of reaction mixture simultaneously in cases and control in triplicate on a StepOnePlus Real-Time PCR System (Applied Biosystems) with SYBR Premix Ex Taq Tli RNaseH Plus (Takara Biotechnology, Dalian, China) and default setting of the reaction condition. The copy number in each sample was determined by using the ΔΔCt method, which compared the ΔCt (cycle threshold) of the target and a copy number neutral region [an ultra-conserved region (https://ccg.epfl.ch/UCNEbase/view.php?data=ucne&entry=5530)] in the case with that of the control. Two independent primer-pairs were used for each validation (Dong et al., 2018a).

## Results

Overall, we recruited 50 pregnancies with fetus with increased NT in the first-trimester Down syndrome screening. 37 CVS samples were collected from the first-trimester and AF samples were collected in later gestational weeks in the other 13 cases. Thirty-four cases were reported to have insolated increased NT with/without other soft markers (68.0%) and 16 cases were diagnosed with syndromic abnormalities (increased NT and fetal structural malformations, Table 1). All cases have undergone routine prenatal diagnosis by CMA and 90% of these cases also have G-banded chromosome results available. CMA with/without karyotyping yielded diagnoses in eight cases, with pathogenic/likely pathogenic (P/LP) CNVs, providing a diagnostic yield of 16.0% (Figure 1 and Table 1).

**Table 1.**
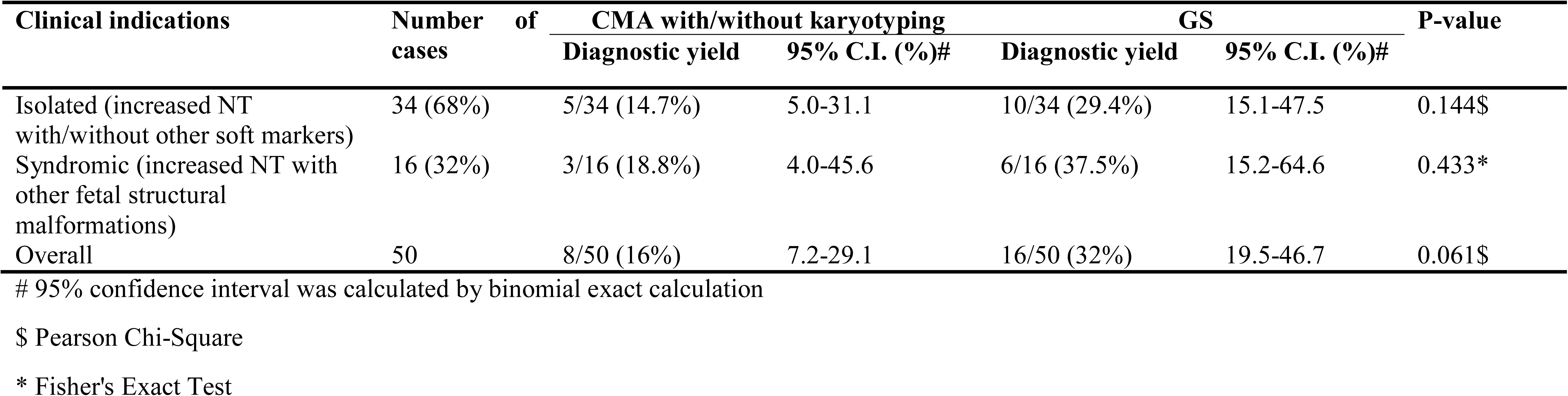
Prenatal detection rates of the fetuses with increased NT by CMA/Karyotyping and GS.

### Numerical Disorders and CNV

In this study, karyotyping identified four microscopic deletions/duplications, while CMA reported eight cases with P/LP CNVs and seven with variants of uncertain significant (VOUS or VUS, **Supplementary Table S2**). Compared with CMA results, GS not only detected all P/LP CNVs and VOUS detectable by CMA, but also defined additional findings including a case of mosaic turner syndrome, a case of comprehensive delineation of CNV in one case with 16p11.2 recurrent deletion syndrome (Table 2) and two additional VOUS (**Supplementary Table S2**).

In 17NT0005, a fetus at the 12^th^ gestational week with NT >14.3-mm and presented with cystic hygroma. Normal CMA results was reported by the SNP platform and no karyotyping results were available (Figure 2A, B and **Supplementary Figure S1**). GS reported a mosaic turner syndrome with mosaic level around 40% (Figure 2C). Further validation by aCGH-based platform confirmed this finding and with consistent mosaic level estimated (Figure 2D). Mosaic turner syndrome was known to be the causative finding for the fetus with cystic hygroma, but it was missed by the original CMA in this study (Alpman et al., 2009). In the original CMA result, no decrease of copy-ratio of the Y chromosome was indicated by the allele difference data (copy-ratio as around 1, Figure 2A), although it was slightly decreased compared with the one of the X chromosome based on the coverage difference data (Figure 2B). It indicated that SNP based platform might not be sensitive enough for detection of mosaicism on chromosome Y. Since the DNA source of this sample was from CVS, we could not exclude the possibility of this mosaic turner syndrome was due to confined placental mosaicism. However, further confirmation by testing with AF sample was not possible as the pregnancy was terminated due to the severe presentation in the fetus (cystic hygroma).

**Table 2.**
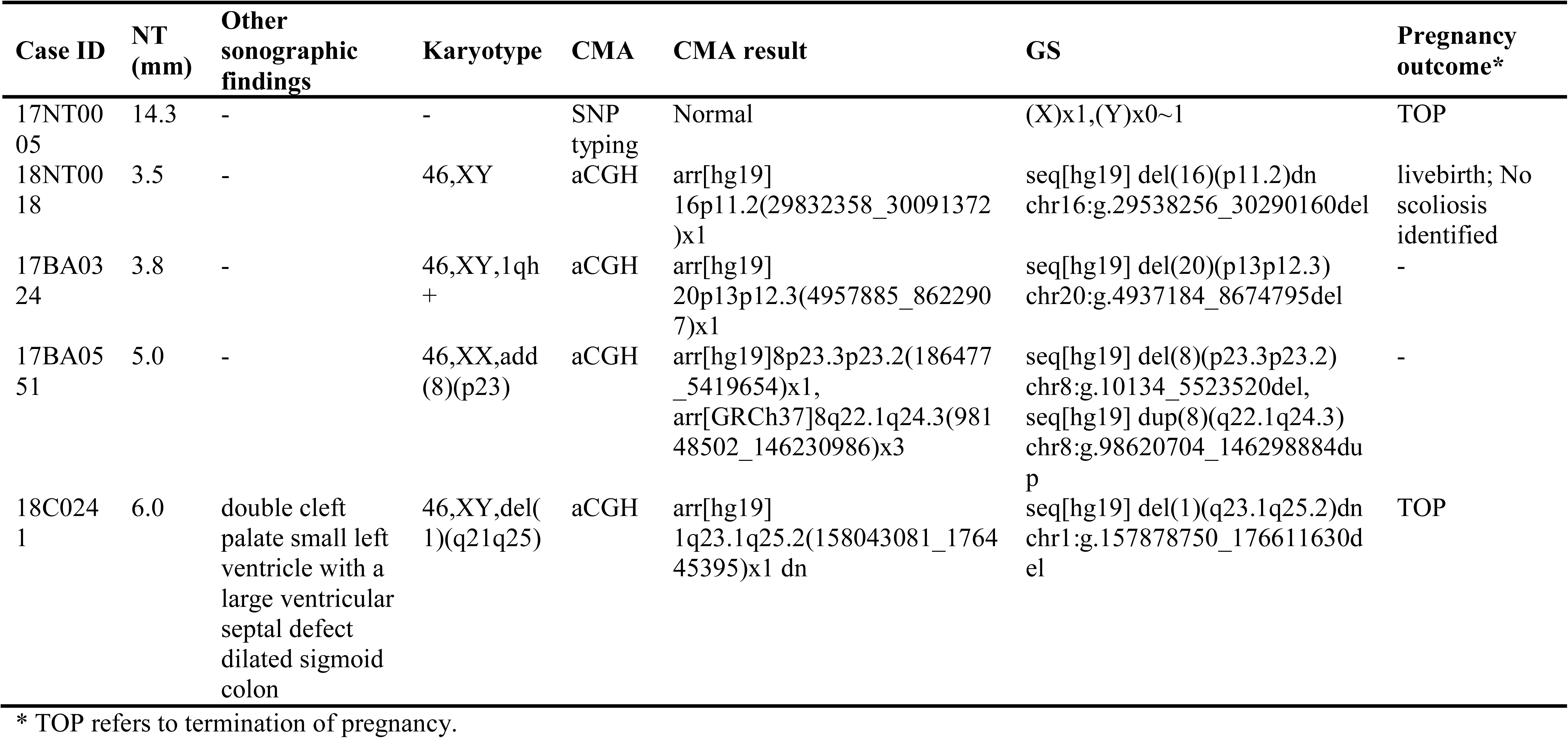
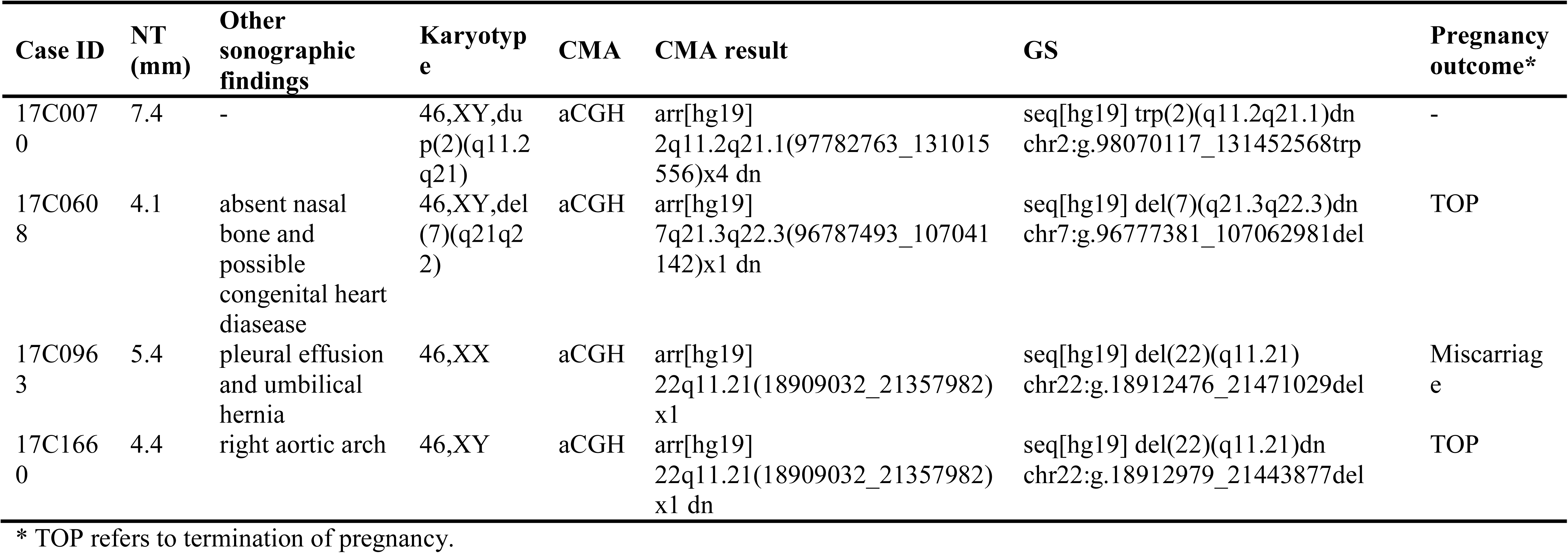
Summary of numerical disorder and pathogenic or likely pathogenic CNVs detected.

| Case ID | NT (mm) | Other sonographic findings | Karyotype | CMA | CMA result | GS | Pregnancy outcome* |
| --- | --- | --- | --- | --- | --- | --- | --- |
| 17NT0005 | 14.3 | - | - | SNP typing | Normal | (X)x1,(Y)x0~1 | TOP |
| 18NT0018 | 3.5 | - | 46,XY | aCGH | arr[hg19] 16p11.2(29832358_30091372)x1 | seq[hg19] del(16)(p11.2)dn chr16:g.29538256_30290160del | livebirth; No scoliosis identified |
| 17BA0324 | 3.8 | - | 46,XY,1qh+ | aCGH | arr[hg19] 20p13p12.3(4957885_8622907)x1 | seq[hg19] del(20)(p13p12.3) chr20:g.4937184_8674795del | - |
| 17BA0551 | 5.0 | - | 46,XX,add(8)(p23) | aCGH | arr[hg19]8p23.3p23.2(186477_5419654)x1, arr[GRCh37]8q22.1q24.3(98148502_146230986)x3 | seq[hg19] del(8)(p23.3p23.2) chr8:g.10134_5523520del, seq[hg19] dup(8)(q22.1q24.3) chr8:g.98620704_146298884dup | - |
| 18C0241 | 6.0 | double cleft palate small left ventricle with a large ventricular septal defect dilated sigmoid colon | 46,XY,del(1)(q21q25) | aCGH | arr[hg19] 1q23.1q25.2(158043081_176445395)x1 dn | seq[hg19] del(1)(q23.1q25.2)dn chr1:g.157878750_176611630del | TOP |
| 17C0070 | 7.4 | - | 46,XY,dup(2)(q11.2q21) | aCGH | arr[hg19] 2q11.2q21.1(97782763_131015556)x4 dn | seq[hg19] trp(2)(q11.2q21.1)dn chr2:g.98070117_131452568trp | - |
| 17C0608 | 4.1 | absent nasal bone and possible congenital heart disease | 46,XY,del(7)(q21q22) | aCGH | arr[hg19] 7q21.3q22.3(96787493_107041142)x1 dn | seq[hg19] del(7)(q21.3q22.3)dn chr7:g.96777381_107062981del | TOP |
| 17C0963 | 5.4 | pleural effusion and umbilical hernia | 46,XX | aCGH | arr[hg19] 22q11.21(18909032_21357982)x1 | seq[hg19] del(22)(q11.21) chr22:g.18912476_21471029del | Miscarriage |
| 17C1660 | 4.4 | right aortic arch | 46,XY | aCGH | arr[hg19] 22q11.21(18909032_21357982)x1 dn | seq[hg19] del(22)(q11.21)dn chr22:g.18912979_21443877del | TOP |

\* TOP refers to termination of pregnancy.

**Figure 2.**
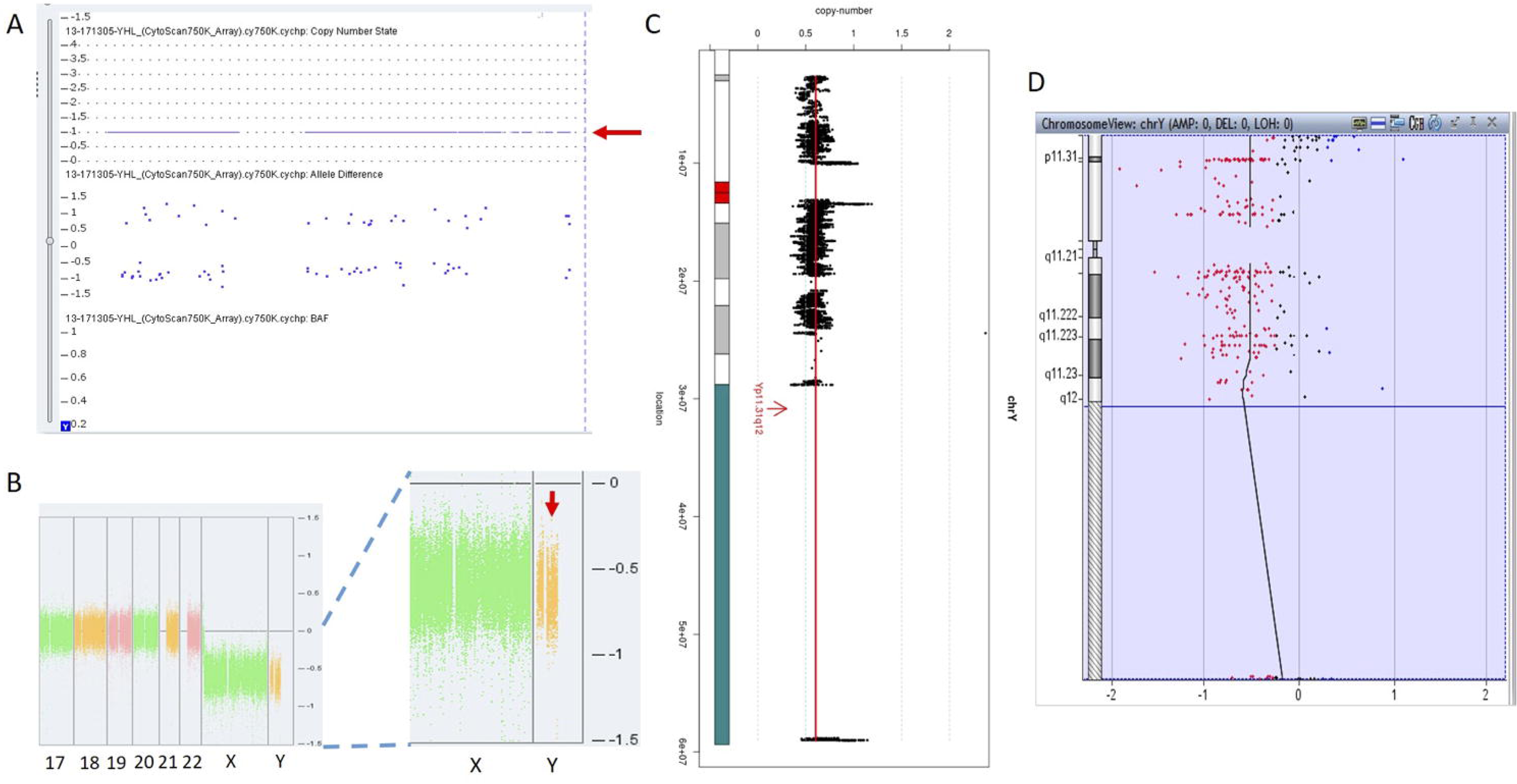
Mosaic Turner Syndrome Detected by GS. In 17NT0005, CytoScan 750K CMA platform reported (**A**) a copy-number as 1 for chromosome Y (indicated by a red arrow) and (**B**) apparently normal male fetus (partial figure of weighted Log2 ratio across each chromosome), although slight decrease of Log2 ratio of chromosome Y was observed compared with the Log2 ratio of chromosome X (manifested by a red arrow). (**C**) GS reported a 40% decrease of the copy-number of chromosome Y and confirmed by the aCGH CMA platform shown in (**D**). In figure (**C**), copy-ratio of each window is indicated by a black dot and the average copy-ratio of chromosome Y is reflected by a red vertical line. Karyogram of chromosome Y is shown in the left. In figure (**D**), dots in red, in black and in blue indicate copy-number lost, copy-number neutral and copy-number gained, respectively. The average Log2 ratio is indicated by a black vertical line.

In total, GS identified one mosaic turner syndrome and seven cases with P/LP microdeletions/microduplications (Table 2) in this group.

### SNV Detection and Interpretation

To demonstrate the ability of detecting SNVs and InDels by GS, we reported seven cases with P/LP point mutations by GS following the ACMG guidelines (Table 3).

**Table 3.** Summary of pathogenic or likely pathogenic mutations revealed by GS.

| Case | NT (mm) | Other sonographic findings | Gene | Mutation and het/hom* | Consequence | Inheritance mode | Disease association(s) [MIM #] | Pregnancy outcome | Inheritance confirmation<sup>#</sup> |
| --- | --- | --- | --- | --- | --- | --- | --- | --- | --- |
| 14C1232 | 3.73 | mild ventricular disproportion | *ARMC4* | NM_018076:c.1614_1615del (p.P538fs) het | Frameshift | AR | Ciliary dyskinesia, primary, 23 (615451) | Live birth | Pat |
|  |  |  |  | NM_018076:c.C2306A (p.P769H) het | Missense | AR | Ciliary dyskinesia, primary, 23 (615451) | Live birth | Mat |
| 15C0337 | 4.18 | - | *ANKRD11* | NM_001256182:c.2404dupC (p.L802fs) het | Frameshift | AD | KBG syndrome (148050) | Live birth; newborn exam: low set ears, increased nuchal fold; right hand single transverse crease; bilateral clinodactyly; postnatal follow-up at three years old showed bilateral hearing loss (40db) | *de novo* |
| 15C0802 | 5.03 | - | *GATA4* | NM_002052:c.C1325T(p.A442V) het | Missense | AD | Atrial septal defect 2 (607941); Tetralogy of Fallot (187500) | Live birth | Mat |
| 18NT019 | 3.5 | - | *NSD1* | NM_022455:c.3797-2A>G het | Splicing-site | AD | Sotos syndrome 1(117550) | - | - |

| Case | NT (m m) | Other sonographic findings | Gene | Mutation and het/hom | Consequence | Inheritance mode | Disease association(s) [MIM #] | Pregnancy outcome | Inheritance confirmation |
| --- | --- | --- | --- | --- | --- | --- | --- | --- | --- |
| 16C1 953 | 10.9 | Large jugular lymphatic sac; cystic hygroma; absent ductus venosus a-wave; bilateral hand with bifid thumb. | *BRAF* | NM_004333: c.G1411T (p.V471F) het | Missense | AD | Noonan syndrome 7 (613706); Cardiofaciocutaneous syndrome (115150) | TOP | - |
| 18NT 0003 | 12 | Hydrosarca; cystic hygroma; short long bones | *COL2A1* | NM_001844: c.G2950A (p.G984S) het | Missense | AD | Achondrogenesis, type II (200610) | TOP | *de novo* |
| 18NT 018 | 3.5 | Reverse 'a' wave in DV, absent nasal bone and possible congenital heart disease: single atrium, single ventricle, bilateral renal volume increased with polycystic changes | *KMT2D* | NM_003482: c.16474delG (p.D5492fs) het | Frameshift | AD | Kabuki syndrome 1(147920) | TOP | - |

\* het/hom refer to heterozygous mutation or homozygous

The first line of our detection was to report the cases with known disease-causing mutations. In case 16C1953, a male fetus presented with increased NT (10.9-mm), cystic hygroma and bilateral large jugular lymphatic sacs, absence of ductus venosus a-wave and bilateral bifid thumbs in the gestational week of 12, was reported with normal CMA and karyotyping. GS detected a pathogenic heterozygous point mutation NM_004333:c.G1411T(p.V471F) in *BRAF* gene, which was reported to cause Cardio-Facio-Cutaneous Syndrome (Abe et al., 2012) and Noonan Syndrome (Nystrom et al., 2008;Croonen et al., 2013) in an autosomal dominant manner (Table 3). While, in 18NT0003, a female fetus with increased NT, hydrosarca and short limbs at the 12^th^ week of gestation, was reported with normal CMA and karyotyping findings. A heterozygous point mutation NM_001844.4:c.G2950A(p.G984S) was detected in an autosomal dominant disease-causing gene *COL2A1* by GS. Since a different base change NM_001844.4:c.G2950C(p.G984R) in the same location was reported to cause Achondrogenesis, type II (OMIM: #200610) or Type II Collagenopathies (Barat-Houari et al., 2016), in addition, the variant was confirmed to be a *de novo* mutation (Figure 3A), thus it was classified as a likely pathogenic causative variant for the fetal phenotype (Table 3).

**Figure 3.**
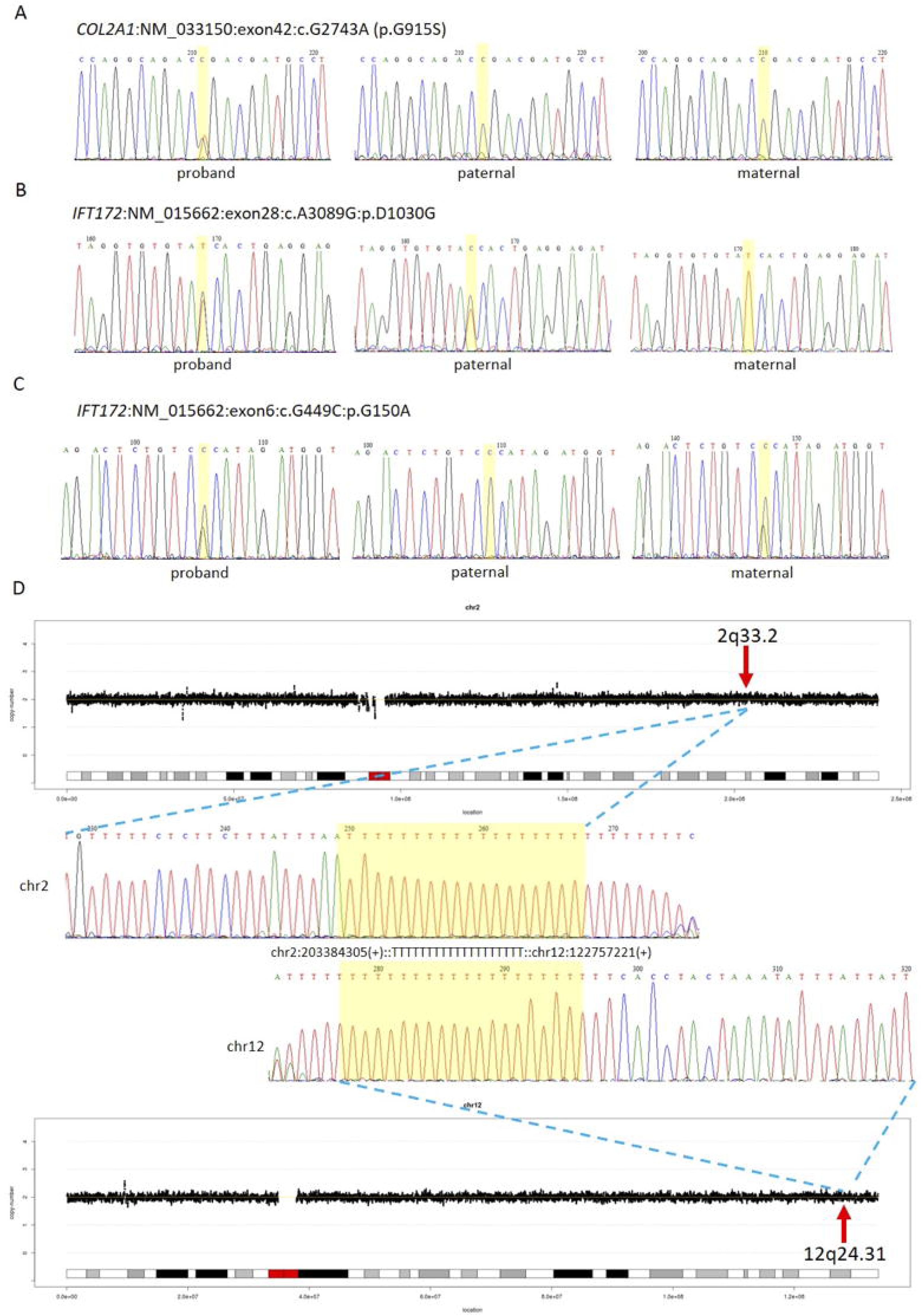
Comprehensive Definition of the Genetic Etiologies in 18NT0003. GS with parental confirmation reported (**A**) a *de novo* heterozygous mutation NM_033150:c.G2743A(p.G915S) in *COL2A1*, (**B**) a paternal heterozygous mutation NM_015662:c.A3089G:p.D1030G in *IFT172* and (**C**) a maternal heterozygous mutation NM_015662:c.G449C:p.G150A in *IFT172*. All the mutated sites are highlighted in yellow. (**D**) Distributions of copy-ratios in chromosome 2 and chromosome 12 are shown in the top and the bottom, respectively. The inserted site in chromosome 2 and the rearranged segments in chromosome 12 are indicated by red arrows with the band numbers, both of these two regions were copy-number neutral. Dots in black indicate the copy-ratio in each window across different chromosomes. X axis and Y axis represent the genomic coordinates and the copy-ratios. Dotted lines in grey indicate the copy-number 0, 1 and 3 and 4, respectively. Sanger sequencing results of the breakpoints are shown in the middle. Forward sequencing is shown in the top, while reverse complementary sequencing result of the reverse sequencing is shown in the bottom. The breakpoint coordinates in chromosome 2 and chromosome 12 are shown with the aligned orientation, while the inserted sequence is remarked in between and highlighted in yellow in both Sanger sequencing results.

In addition, we also identified five novel mutations in four samples, including variants in three autosomal dominant and one autosomal recessive genes (Table 3). For instance, 18NT018, a female fetus with increased NT, absent nasal bone and reverse a-wave ductus venosus observed in the 1^st^ trimester ultrasound screening with both normal karyotyping and CMA. Morphology scan in the 2^nd^ trimester revealed complex congenital heart disease (single atrium and ventricle) and polycystic kidneys. GS identified a novel heterozygous deletion NM_003482:c.16474delG(p.D5492fs) which resulted in a frameshift in the *KMT2D* gene. Mutations in *KMT2D* have been known to cause Kabuki syndrome (OMIM: #602113) in an autosomal dominant manner, commonly resulting in heart defects and renal malformations consistent with the presentation of this fetus. Therefore, it was further classified as likely pathogenic (Table 3). Compound heterozygous point mutations were found in the fetus 14C1232: a paternally inherited frameshift deletion NM_018076:c.1614_1615del(p.P538fs) and a maternally inherited nonsynonymous mutation NM_018076:c.C2306A(p.P769H) were detected by GS and further confirmed by Sanger sequencing (**Supplementary Figure S2**). Mutations in gene *ARMC4* in an autosomal recessive manner are known to cause ciliopathies such as ciliary dyskinesia, primary, 23 (OMIM: # 615451). The couple decided to keep the pregnancy and resulted in a live born. Follow-up study is on-going.

Furthermore, GS also detected 23 VOUS in 18 cases (Figure 3B and **C**, **Supplementary Table S3**).

### SV Detection and the Potential Pathogenicity

Apart from detecting CNV and point mutations, GS was able to identify structural rearrangements by utilizing paired-end reads. Overall, two inversions and two insertions were detected among these 50 cases (**Supplementary Table S4**).

In 18NT0003 described above, GS also reported a cryptic balanced complex insertion seq[hg19]ins(2;12)(q33.2;q24.31)g.[chr2:203384219_203384293inschr12:122757221_122907271cx]chr12:g.122757221_122907271del carrying a partial segment of gene *CLIP1* from chromosome 12 to the site of *BMPR2* in chromosome 2 (**Supplementary Table S4**, Figure 3D). The original fragment from chromosome 12 (seq[hg19] chr12:122757221_122907271) was 150.1-kb in size but it was divided into 11 fragments with five segments lost when inserted to chromosome 2 (**Supplementary Figure S3**). There were 18 continuing nucleotide Ts found in the breakpoint junction of the inserted sit, not belonging to either chromosomes (2 and 12) (Figure 3D), indicating that the breakpoint repairing mechanism might be caused to non-homologous end joining with a cryptic insertion (Carvalho and Lupski, 2016). Parental confirmation by PCR and Sanger sequencing established this insertion to be a paternally inherited event. Regarding the potential pathogenicity, it has been known that mutations leading to loss of function in *BMPR2* would cause Primary Pulmonary Hypertension 1 and/or Pulmonary Venooclusive Disease 1 (OMIM: #600799). However, we were unable to determine whether the disruption of *BMRP2* was one of the causal factors of the fetal phenotype, since the pregnancy was terminated. We further followed up with the father (31-yo) and found sinus bradycardia by routine electrocardiograph. Although pulmonary hypertension may be later onset, sinus bradycardia might be one of the markers indicating pulmonary hypertension (Rajdev et al., 2012). Further follow-up is also on-going.

For the other cryptic insertion and two inversions **(Supplementary Table S4)**, since no OMIM genes were involved and no topological associated domains were disrupted, they were further classified as polymorphisms.

### Overall Diagnostic Yield for Fetuses with Increased NT by GS

For these 50 cases, 68.0% of these cases had insolated increased NT with/without other soft markers. However, no significant difference of the diagnostic yields was found between isolated and syndromic groups (Table 1, Chi-Square test P=0.5674). In addition, GS provided a 2-fold diagnostic yield in all 50 cases compared with the routine test by CMA and/or karyotyping.

### Comprehensive Detection and Delineation of the Variants in Individuals

In addition to the identification of individual variants (i.e., SNVs, CNVs and SVs), another advantage of using GS is to delineate the mutated regions involving various types of aberrations. For example, in 18NT0018, a male fetus with isolated increased NT, CMA reported a 259.0-kb *de novo* heterozygous deletion arr[hg19] 16p11.2(29832358_30091372)x1, which was diagnosed as 16p11.2 recurrent deletion syndrome (Lin et al., 2018). While GS refined the breakpoints of the CNV to a 751.9-kb *de novo* deletion seq[hg19] del(16p11.2) chr16:g.29538256_30290160del (Figure 4A) which involved the *TBX6* gene (T-Box 6, Figure 4B) not covered by the CNV reported by CMA. Heterozygous deletion of *TBX6* was further confirmed by qPCR experiment (Figure 4C) in the proband and the parents. The smaller CNV reported by CMA was contributed by the absence of probes in the region next to the reported region (Figure 4B). *TBX6* is currently involved in 16p11.2 recurrent deletion syndrome (Lin et al., 2018) and is highly correlated with scoliosis if there was a presence of a hemizygous T-C-A haplotype (Wu et al., 2015;Liu et al., 2019). However, there was no mutant allele detected for each of these three common single-nucleotide polymorphisms (SNPs: rs2289292, rs3809624 and rs3809627) in the fetus. Further follow-up with the family: the pregnancy was kept and no scoliosis was found in the infant, echoing lower risk of having scoliosis in the absence of a hemizygous T-C-A haplotype in patient with 16p11.2 recurrent deletion syndrome (Wu et al., 2015;Liu et al., 2019).

**Figure 4.**
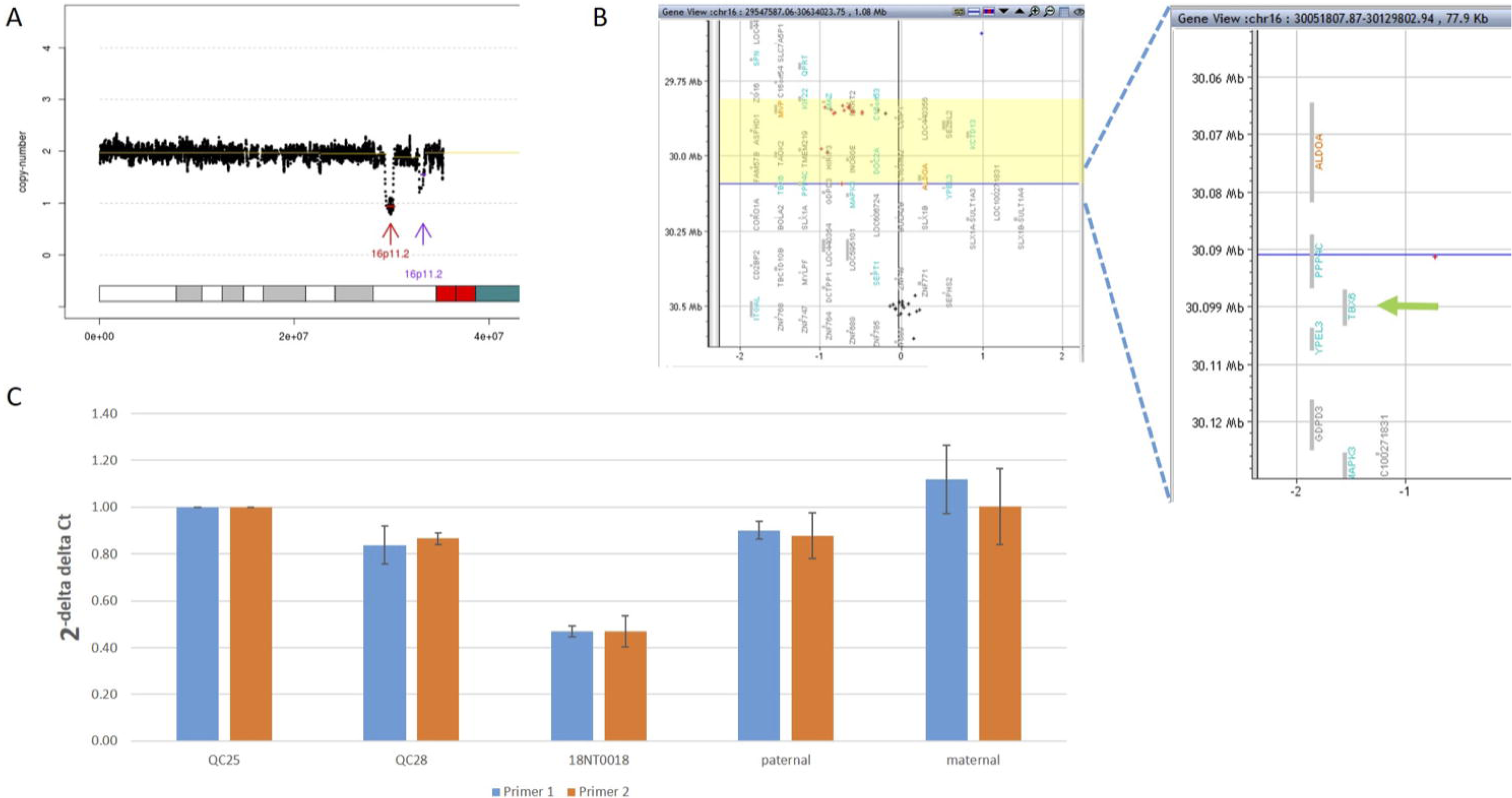
Comprehensive Delineation of 16p11.2 Recurrent Deletion Syndrome. In 18NT0018, (**A**) GS reported a 751.9-kb *de novo* pathogenic deletion seq[hg19] del(16)(p11.2) chr16:g.29538256_30290160del (indicated by a red arrow) and a polymorphism deletion (population-based U-test P>0.01, indicated by a purple arrow) in chromosome 16p11.2; (**B**) original CMA reported a 259.0-kb heterozygous deletion arr[hg19] 16p11.2(29832358_30091372)x1 highlighted in yellow and without a probe locating in gene *TBX6*. (**C**) quantitative PCR two independent pairs of primers targeting *TBX6* shows approximately 0.5 copy-ratio in 18NT0018 compared with normal control (Q25). The parental experiments confirmed the heterozygous deletion in 18NT0018 is in *de novo* manner.

## Discussion

In this study, we applied GS for 50 fetuses with increased NT with/without other fetal structural malformations. In comparison, GS provided an overall diagnostic rate of 32.0%, which was a 2-fold increase compared to the current prenatal diagnosis tests (16.0%). Additional diagnoses include one mosaic turner syndrome, eight cases with P/LP CNVs (Table 2) and seven cases with P/LP point mutations (Table 3). In addition, GS reported two cryptic insertions and two inversions and we further demonstrated the potential pathogenicity of disrupting OMIM disease-causing gene by the inserted site in the follow-up study. Overall, GS demonstrates the ability to detect the various disease-causing variants in human diseases such as fetuses with increased NT (Figure 5).

**Figure 5.**
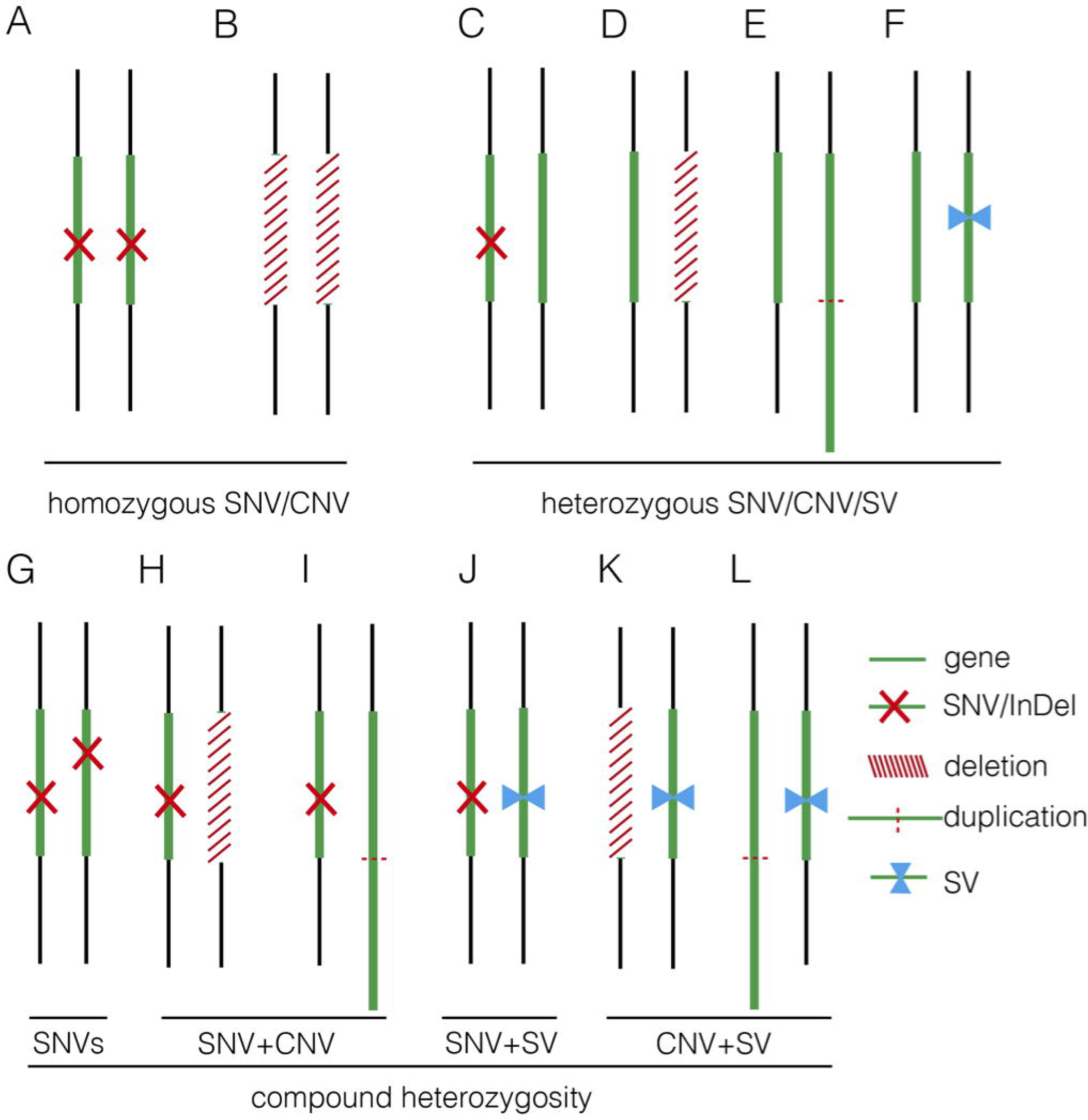
Summary of Various Genomic Variants Proposed to Contribute to Human Diseases. Figures (A) and (B) shows the disease was caused by a homozygous SNV or a homozygous deletion commonly affecting a disease-causing gene in autosomal recessive manner; Figures (C-F) indicate the disease was caused by a heterozygous SNV [NM_004333: c.G1411T (p.V471F) in 16C1953], a heterozygous deletion (seq[hg19] del(16)(p11.2)dn chr16:g.29538256_30290160del in 18NT0018), a triplication (seq[hg19] trp(2)(q11.2q21.1)dn chr2:g.98070117_131452568trp in 17C0070) and a heterozygous SV (seq[hg19]ins(2;12)(q33.2;q24.31)g.[chr2:203384219_203384293inschr12:122757221_1 22907271cx] chr12:g.122757221_122907271del in 18NT0003), respectively, commonly affecting a disease-causing gene in autosomal dominant manner; Figures (G-L) show the disease was contributed by compound heterozygosity a combination of two or more variants. Figure G shows the typical mode of compound heterozygosity (NM_018076:c.1614_1615del (p.P538fs) and NM_018076: c.C2306A (p.P769H) from different parental origin in 14C1232). A gene content is indicated by a green bar. SNV/InDel, deletion, duplication and SV are indicated by a red cross, a white bar (with red slashes), elongated green bar (with red dotted line), and a pair of opposite blue arrows, respectively.

For the detection of numerical disorders and CNVs, GS detected additional findings including one case of mosaic turner syndrome and one case of comprehensive delineation of 16p11.2 recurrent deletion syndrome, compared with CMA (Table 2). For the case with the mosaic turner syndrome (∼40% mosaic level) additionally detected by GS (Figure 2), it was undetected by CMA possibly due to the algorithm employed for calling the copy-number changes by calculating the allele differences. Further confirmation of this mosaic finding by using aCGH-based CMA platform shows the cross platform differences of CMA testing (Haraksingh et al., 2017;Wang et al., 2019). Of noted, in routine setting, QF-PCR experiment was conducted prior to CMA for the detection of MCC and common aneuploidy. However, the missed detection of decreased copy-ratio of the Y chromosome by QF-PCR in this case might be due to the low level mosaicism presented (Shin et al., 2016). In addition, GS was able to delineate the 16p11.2 recurrent deletion which included the disease-causing gene *TBX6*, missed detection of which in this customized CMA platform was owing to the lack of probes located in the target region (Figure 4B). In this study, the detection results from only two CMA platforms were used for comparison and the missed detections and delineations described might be limited to these specific CMA platforms. A large scale and systematic comparison of the performance of different currently available CMA platforms in prenatal diagnosis is warranted as certain platforms such as CytoScan HD and CytoSNP 850K have demonstrated to provide accurate breakpoints of CNVs detected and reliable detection of mosaicism, level of which greater than 20-30% (Haraksingh et al., 2017;Wang et al., 2019).

On top of the scope of current prenatal genetic diagnosis by karyotyping and CMA, GS also detected seven cases with P/LP SNVs and InDels, including three known pathogenic mutations/sites and five novel mutations (in four cases). The incidence of P/LP mutations in this study was 14.0%, which was higher than the newly reported studies utilizing trio-based WES (Lord et al., 2019;Petrovski et al., 2019). A possible reason is the limited sample size or echoing the message that GS is more powerful than WES for detecting exome variants, particularly for SNVs and InDels (Belkadi et al., 2015). However, no WES data was available for comparison in our study.

Current clinical guidelines for variants interpretation are available for SNVs/InDels and CNVs, but there are no guidelines available for SV interpretation. Our study demonstrates the feasibility of detecting SVs by utilizing paired-end reads from GS. With that, we were able to identify inversions and insertions and to show the potential pathogenicity in a paternally inherited insertion disruption of gene *BMRP2*. Follow-up study conducted in the father indicated a sinus bradycardia from routine electrocardiograph. Detection of such cryptic rearrangements would help us explore a “blind spot” by the routine methods. However, CNVs and SVs are predominantly mediated by repetitive elements [commonly >1-kb (van Heesch et al., 2013)], which causes difficulty in the identification of the flanking unique sequences by standard GS with small-insert libraries employed in this study (Chen et al., 2008;Carvalho and Lupski, 2016;Dong et al., 2018a). Further study with the other approaches such as mate-pair sequencing might be an alternative method for identifying additional pathogenic SVs (Talkowski et al., 2012a;Zirui et al., 2018), which is enriched in the disease groups (Talkowski et al., 2012b). Overall, apart from identifying individual genomic variants, GS also shows its advantages in comprehensively defining different mutation types in different alleles. It is supported by the absence of a hemizygous T-C-A haplotype in the region of 16p11.2 recurrent deletion syndrome (Wu et al., 2015;Liu et al., 2019) (Figure 5).

Overall, by taking together the various genomic variants, the diagnostic rates of GS were not significantly different between the isolated and syndromic groups (Table 1). For the isolated group, GS provided a diagnostic yield as 29.4%, which was significantly higher than the previously reported (Lord et al., 2019). One of the reasons might be owing to the small sample size. Nonetheless, our current data truly show a higher diagnostic yield. Further study with larger sample size is warranted. However, there were still 34 cases without a positive diagnosis by genome sequencing. Pregnancy outcomes were available for 18 cases, seven (38.9%) of which were terminated pregnancies due to severe fetal malformations, such as pleural effusion, multicystic dysplastic kidneys and complex congenital heart disease. It indicates that genome sequencing is still not revealing all the causative variants and the clinical decision of continuing or termination of pregnancy is largely dependent on the severity of fetal malformation(s) found in each particular case. Nonetheless, re-analysis of the GS data frequently upon the updating literatures might be able to increase the genetic diagnosis, thus, provide the causative answers for the family.

In this study, we aimed to evaluate the feasibility of applying GS for prenatal diagnosis in the fetuses with increased NT. Apart from the increased diagnostic yield, the requirement of DNA amount as input for GS (100-ng) was less, compared with CMA (250-ng for CytoScan 750K or 300-ng for 8×60K Fetal Chip), which would further facilitate GS in prenatal diagnosis, particularly for the AF samples from early gestational weeks. In addition, in comparison with the methods for routine prenatal diagnosis (CMA and karyotyping), GS would be able to provide a comparable turn-around-time of 10 working days from DNA preparation to variant validation (Figure 1). Although the reagent cost of GS is less than 1,000 USD per case, the demands of computational resources and labor costs for data analysis and interpretation are still higher than routine methods (Figure 1). Parental confirmation is important for determination of the variants’ pathogenicity. In this study, we applied GS only for the proband and conducted the parental confirmation by Sanger sequencing, which was laborious. Although applying trio-based GS testing would be helpful for variant filtering in initial data analysis, it would triplicate the cost, thus, preventing the value of clinical application.

Regarding the interpretation of genomic variants, similarly with the prenatal studies by WES, even with parental confirmation, GS study would still yield a number of VOUS, the pathogenicity and clinical significance of which are still uncertain. One of the reasons would be lack of clinical symptom(s) in early pregnancy; follow-up with detailed diagnosis would be warranted. For instance, in 18NT0003, a fetus with increased NT, hydrosarca and short limbs, GS reported a known LP mutation in *COL2A1* that might already explain the malformation. In addition, GS with parental confirmation also reported another two variants in compound heterozygosity manner, including a paternally inherited heterozygous variant NM_015662:c.A3089G(p.D1030G) and a maternally inherited heterozygous variant NM_015662:c.G449C(p.G150A) in gene *IFT172* (Figure 3B **and C**). Mutations in *IFT172* are known to cause retinitis pigmentosa (OMIM: #616394) or short-rib thoracic dysplasia (OMIM: # 615630), the latter might be also be partially correlated with the fetal phenotype but might only present in later gestation. However, further confirmation of the pathogenicity by correlating the phenotype in later gestational weeks was not possible as the pregnancy was terminated. Nonetheless, GS still provides a comprehensive foundation for gene discovery. Moreover, one limitation in this study was the retrospective approach and the limited sample size, a prospective back-to-back in comparison of GS’s performance with routine prenatal diagnosis approaches with larger sample size and other clinical indications are warranted in near future.

In conclusion, our study demonstrates that applying GS in fetuses with increased NT can provide a comprehensive detection of various disease-causing genomic variants, including SNVs, InDels, CNVs and structural rearrangements. Compared with current routine prenatal diagnosis approaches, GS not only shows increased sensitivity of detecting mosaic numerical disorders and comprehensive delineation of P/LP CNVs, but also provides the ability of identifying causative point mutations and balanced structural rearrangements that are beyond the detection scope of current prenatal diagnosis protocols. Although future studies with larger sample sizes and in prospective manners are warranted, given comparable turn-around-time and less DNA required in addition to the increased diagnostic yield, our study provides the first and strong evidence to facilitate GS in prenatal diagnosis, particularly in fetuses with increased NT.

## AVAILABILITY

Whole-genome sequencing data used in this study has been made available in the CNGB Nucleotide Sequence Archive (CNSA: https://db.cngb.org/cnsa) under the accession number CNP0000437.

## DISCLOSURE

The authors declare no conflict of interest.

## ACKNOWLEDGMENTS

This project is supported by the National Natural Science Foundation of China (81741004, 81671470 and 31801042), the Health and Medical Research Fund (04152666), Guangzhou Science and Technology Program (2014 Y2-00551, 201504282321393, 201604020078 and 201604020091), and the National Key Research and Development Program of China (2018YFC1004104).

## AUTHOR CONTRIBUTIONS

K.W.C., M.C., T.Y.L. and Z.D. designed the study. H.W., J.C., R.Z., O.Y.M.C., Y.Z. and T.Y.L. collected the samples and followed up. K.W.C., M.S., Z.Y., Y.C., Y.K.K. and Z.D. performed the analysis and data interpretation. M.S., H.Y., Y.W., S.C. and M.H.K.C. conducted the validation. K.W.C., H.W., M.S., J.C. and Z.D. wrote the manuscript.

