## Supplemental for "Prenatal Diagnosis of Fetuses with Increased Nuchal Translucency by Genome Sequencing Analysis"

**Supplementary Materials and Methods**

**Supplementary Figures**

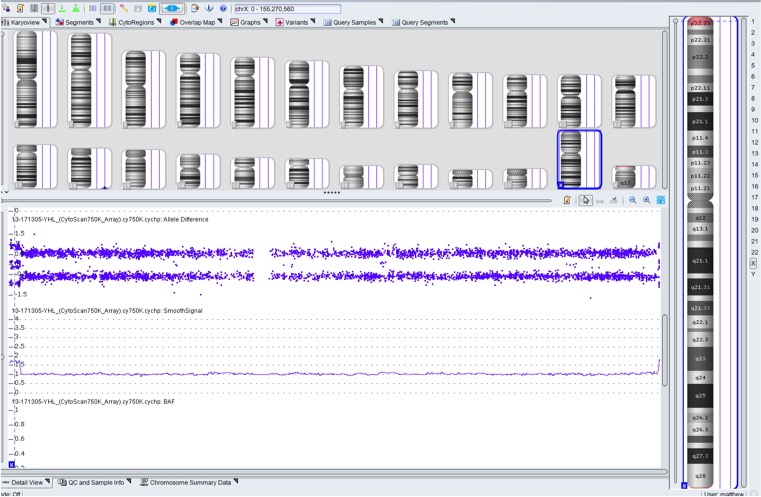

**Supplementary Figure S1. Distribution of copy-ratios in X chromosome in sample** 17NT0005**.** CytoScan 750K chromosomal microarray analysis platform reported copy-number as 1 for chromosome X indicated by a blue line in the middle panel. The figure shows only two genotypes (homozygous single nucleotide polymorphisms, indicated by blue dots in upper panel) presented in this sample reflecting there is only one X chromosome detected.

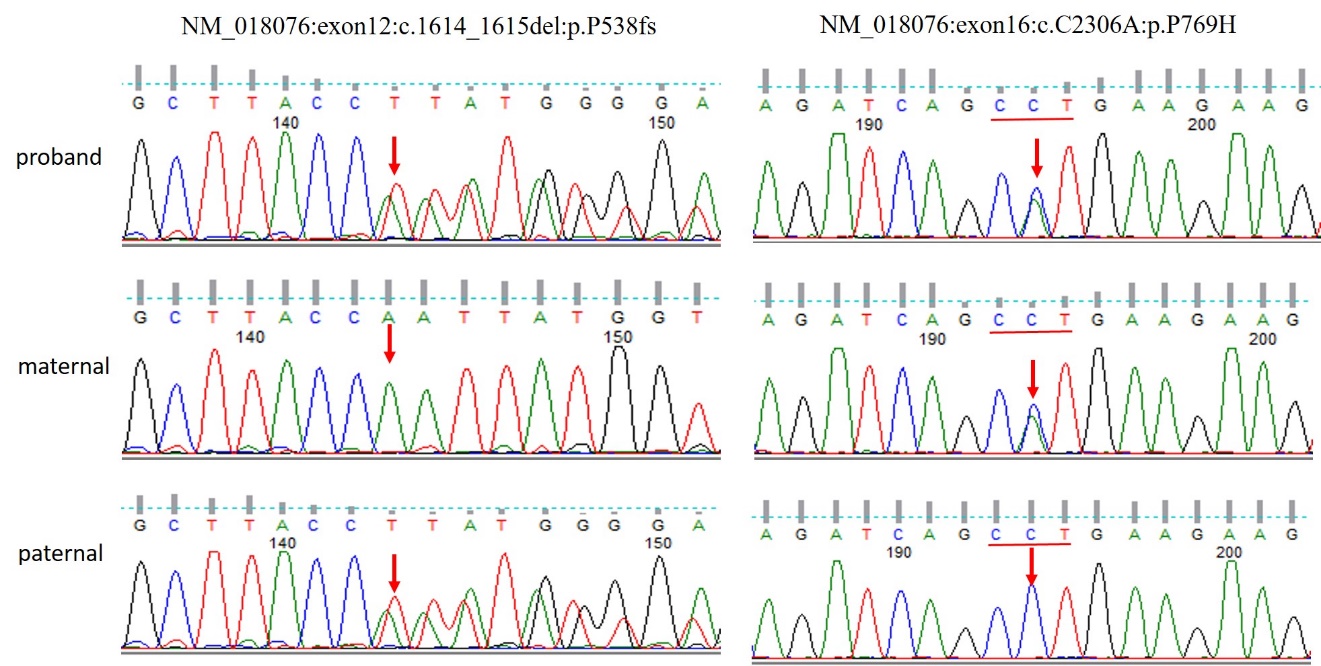

**Supplementary Figure S2. Compound heterozygosity mutations detected in *ARMC4* in sample 14C1232.** Sanger sequencing confirmed a paternally inherited frameshift deletion NM_018076:c.1614_1615del(p.P538fs) and a maternally inherited nonsynonymous mutation NM_018076:c.C2306A(p.P769H) in gene *ARMC4* in fetus 14C1232. The mutated site was highlighted by a red arrow in each figure.

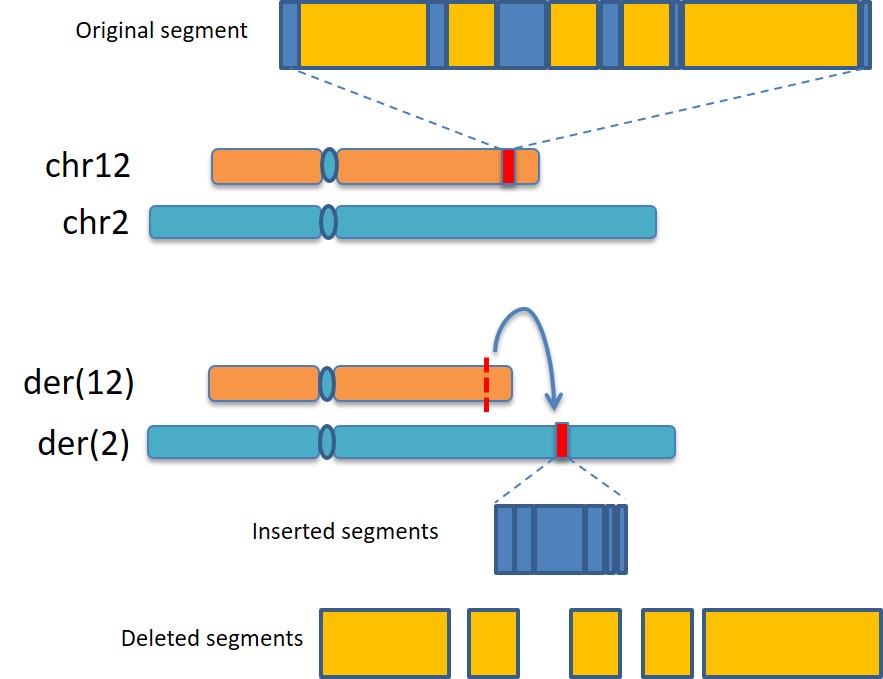

**Supplementary Figure S3. Diagram of paternal inherited insertion detected.** In 18NT0003, fetus with cystic hygroma detected, GS reported a cryptic apparently balanced complex insertion seq[hg19]ins(2;12)(q33.2;q24.31)g.[chr2:203384219_203384293inschr12:122757221_122907271cx]chr12:g.122757221_122907271del. In the upper panel, figure shows the original compositions of chromosomes 12 and 2. The original fragment from chromosome 12 is indicated by a red box with zooming figure showing the composition. In the bottom panel, figure shows the derivative chromosomes 12 and 2 with the fragment from chromosome 12 inserted into chromosome 2 (indicated by a blue arrow). The final composition of the inserted fragment (in blue) and the lost segments (in yellow) are shown below.

**Supplementary Table S1. Primers used in this study**

| No | Validation purpose of the usage of this pair of primers | Forward | Reverse |
| --- | --- | --- | --- |
| 1 | *TBX6* deletion_1 | CCTCATAACATCCGTCACAGTTC | AGGTTATGTCGGACAGTAAAGGT |
| 2 | *TBX6* deletion_2 | GGTCTAAGCCACACACTAACCTA | CTAGAATCTGACTATGGCCTGGG |
| 3 | ins(2;12)-breakpoint-1 | GCCGAGATTGTACAGCCGTA | ATTGACACAACGTAACACCAGC |
| 4 | ins(2;12)-breakpoint-2 | CCACCTGTCCCACTGCAG | GGATGGAAAGGGGAATTTAAACAT |
| 5 | NM_018076:c.1614_1615del (p.P538fs) | ATTCTTAAGGGCAAGCGATACAG | CTTAACTGGGGAAGGATTTTAGGT |
| 6 | NM_018076:c.C2306A (p.P769H) | CACCAACTGCTTTTGTAACATTCA | CTGGGTTAGACGTCAGCTCTAC |
| 7 | NM_001256182:c.2404dupC (p.L802fs) | CAAGTCAGAAAACCACCGATCTC | AAGAAGAAATCAAAAGACCGGCC |
| 8 | NM_002052:c.C1325T(p.A442V) | GTTCTCAGTCAGTGCGATGTC | ACCAGATTGTCGACTTCAAGTTT |
| 9 | NM_004333:c.G1411T (p.V471F) | AGTTTATTGATGCGAACAGTGAA | CCTCTCAGGCATAAGGTAATGTAC |
| 10 | NM_001844:c.G2950A (p.G984S) | AAGGATGATGAGTGCAAGTGGT | ACAGAGCATGGGGTAGGAG |
| 11 | NM_003482:c.16474delG (p.D5492fs) | GGTTAGCTGAAAAGAGAAGAGGG | GGATTTAGTGGTCCCTGAGAGT |
| 12 | NM_022455:c.3797-2A>G | CCCCAGCTCCTTTTGCATAT | CCACTAAATCCTAAGTACTGTGCTAG |
| 13 | NM_000548: c.C4349G (p.P1450R) | CTCCCGAGCTGCAGACTC | CTGTGGCTCTGCTCTTTAAGG |
| 14 | NM_001142519: c.1598_1599del (p.D533fs) | GAATTACTCCTGTGCCACTTAGT | TTGCTTAATATCAAGGAGGATGGA |
| 15 | NM_000347:c.G3958A (p.A1320T) | TGCCACGTTCTGTACTGTCA | GATAGTCAGCCCACTCTTTATGC |
| 16 | NM_003797:c.A841C (p.N281H) | TGGCCTATTTAGATGAAACCCAA | TGCTACTAAGTTAAGATGACAACACA |
| 17 | NM_004380:c.C2399T (p.P800L) | GCATTCTGGAATTTTAATTTCCACGA | GTTGTGAAGCCTTGGATCATTCT |
| 18 | NM_170784:c.G459C (p.L153F) | TCCTAAAATGATGTGGCCTTCAG | GACAGCCTCCATACAGAATCATG |
| 19 | NM_170784:c.C170T (p.T57I) | AATCACTGAAGCTTGACACATGA | GGTCTTCACTAGTTACCACCTGA |
| 20 | NM_022068:c.A6946G (p.T2316A) | ATTCCCACTGCTGAATTCATCCT | TGATGAAGGGGAAATTCTGTTGG |
| 21 | NM_006767:c.C1396T (p.R466W) | CTGGGCAGTGGGAATTTCG | TGATCTTGTCGGTGTAGAGGA |
| 22 | NM_015662:c.A3089G(p.D1030G) | GTACATGTTCACTGTTGCCTTCC | GATGGGGTCTGAGGATGTGTAC |
| 23 | NM_015662:c.G449C(p.G150A) | TCCTGCCTTTCATACTATGGTCT | TGGTAGGGATAATGGTTCTGGAG |

**Continued supplementary Table S1. Primers used in this study**

| No | Validation purpose of the usage of this pair of primers | Forward | Reverse |
| --- | --- | --- | --- |
| 24 | NM_004247:c.T1300A(p.C434S) | CAGAAATCAGTATCCCCACACAG | GTGACTCCAAATCTTCTAGCAGC |
| 25 | NM_024411:c.G638A (p.R213H) | GAGGATGAATGAATGCACTCCAA | GACTGGCACACTCTATCTCGC |
| 26 | NM_001029882:c.C1417T (p.R473C) | CATCTTCAGCAGCGGGGT | AAAGATCCTGTGTCGCCGG |
| 27 | NM_001844: c.C3296T(p.P1099L) | GAACAAGACAGACACCGATTGAG | TGAATTCAGTAAAAGCCGCCTTC |
| 28 | NM_006767:c.T2387C (p.I796T) | AGAACCTGGAGATGAACGTGAC | TCCCAAGGATGAAGCAAGGT |
| 29 | NM_000214:c.C3286T(p.R1096W) | GAGTTCTTGTTCTCATAATCCTTGAT | AGTTGTGAAGTGACCATATGCAA |
| 30 | NM_015559:c.C1387T(p.R463C) | ACTCAAGTCATGTCCGGATTACTA | TCAGTTTATCAGAGGTGGCCATA |
| 31 | NM_016952:c.C2654A(p.S885X) | TAGTCAGAAGCAGTAATCCAGGG | TGCCGTGTAATAGAAATGCTGTT |
| 32 | NM_023110:c.T20G(p.L7R) | ACATCACCTGCAACCATATCA | ACAATTTTCTTCTGGACGCCT |
| 33 | NM_032888:c.C2159T(p.P720L) | CTGATGTCCTCTGGCCTAAGAA | CTCTGGGGCTGGCTCACA |
| 34 | NM_032888:c.C5462T(p.T1821I) | AGGTTTAGTTACAGTGATGTTCATGA | CTCTTCCCTCCTCCGTGAG |
| 35 | NM_000257:c.C5279T(p.T1760M) | CTCATGCCCTTCACCGACT | AGGAGCTGATTGAGACTAGTGA |
| 36 | NM_004004:c.G109A(p.V37I) | GATGTGGGAGATGGGGAAGT | AGAAGATGGATTGGGGCACG |
| 37 | NM_004004:c.235delC(p.L79fs) | TCTTCTTCTCATGTCTCCGGT | AAAGGAGGTGTGGGGAGATG |

**Supplementary Table S2. Variants of unknown significance in copy-number variants detected by CMA and GS**

| Case ID | NT (mm) | Other sonographic findings or clinical indication(s) | Karyotype | CMA platform | CMA result | GS |
| --- | --- | --- | --- | --- | --- | --- |
| 18NT018 | 3.5 | reverse 'a' wave in DV, absent nasal bone and possible congenital heart disease: single atrium, single ventricle, bilateral renal volume increased with polycystic changes | 46,XX | aCGH | Normal | seq[hg19] dup(8)(q24.12) chr8:g.120716217_121150420dup |
| 18NT030 | 3.5 | - | 46,XY | aCGH | Normal | seq[hg19] dup(4)(p16.1) chr4:g.9299672_10258522dup |
| 17C1093 | 3.5 | TTTS stage 3 | 46,XY | aCGH | arr[hg19] 1q21.1(145413384_145642403)x3 | seq[hg19] dup(1)(q21.1) chr1:g.145398417_145638434dup |
| 16C1303 | 3.5 | bilateral jugular lymphatic sac | 46,XY | aCGH | arr[hg19] 4p16.3(1562122_1925699)x3 pat | seq[hg19] dup(4)(p16.3)pat chr4:g.1504940_1920056dup |
| 15C0337 | 4.18 | positive Down screening risk:T21:1:3, T13 & T18 positive | 46,XX | aCGH | arr[hg19] 21q22.11(32914256-33040597)x3 pat | seq[hg19] dup(21)(q22.11)pat chr21:g.32828629_33107471dup |
| 18C0028 | 4.4 | Micrognathia | 46,XX | aCGH | arr[hg19] 17q25.3(79202056_79964653)x1 dn | seq[hg19] del(17)(q25.3)dn chr17:g.79163475_79960318del |
| 18C0096 | 5.3 | positive Down screening risk: 1:2, T13 & T18 postive | 46,XY | aCGH | arr[hg19] 8p23.3(190264_489868)x3 mat | seq[hg19] dup(8)(p23.3)mat chr8:g.10132_557003dup |
| 18C0588 | 3.7- 4.1 | bilateral multicystic dysplastic kidneys; reverse a-wave ductus venosus; ventriculomegaly (10.5-mm) | 46,XX | aCGH | arr[hg19] 6p24.3p24.2(10055082_11018204)x1 pat | seq[hg19] del(6)(p24.3p24.2)pat chr6:g.10068225_11182560del |
| 18C0941 | 5.0 | positive Down screening risk: DSS1: 1:2, T18 and T13 both positive | 46,XX | aCGH | arr[hg19] 15q11.2(22834488_23128364)x1 pat | seq[hg19] del(15)(q11.2)pat chr15:g.22680769_23423791del |

**Supplementary Table S3. Variants of unknown significance in SNVs/InDels detected by GS**

| **Case** | **NT (mm)** | **Other sonographic findings** | **Gene** | **Mutation (all heterozygous)** | **Consequence** | **Inheritance mode** | **Disease association(s) [MIM #]** | **Pregnancy outcome** | **Inheritance confirmation** |
| --- | --- | --- | --- | --- | --- | --- | --- | --- | --- |
| 14C0520 | 4.1 | Absent nasal bone, reverse ductus venosus a-wave | *TSC2* | NM_000548: c.C4349G (p.P1450R) | Missense | AD | Tuberous sclerosis-2 (613254) | - | - |
| 14C1309 | 6.21 | - | *FAM111A* | NM_001142519: c.1598_1599del (p.D533fs) | Frameshift | AD | Gracile bone dysplasia (602361); Kenny-Caffey syndrome, type 2 (127000) | Live birth | Mat |
| 14C1348 | 10.4 | bilateral mild pleural effusion, minimal ascites; bilateral pulmonary hypoplasia; severe edema around heads and whole body; one single umbilical artery | *SPTB* | NM_000347:c.G3958A (p.A1320T) | Missense | NA/AD | Anemia, neonatal hemolytic, fatal or near-fatal; Spherocytosis, type 2 (616649) | TOP | Mat |
| 15C0076 | 7.43 | reverse ductus venosus a-wave | *EED* | NM_003797: c.A841C (p.N281H) | Missense | AD | Cohen-Gibson syndrome (617561) | Live birth | Mat |
|  |  |  | *CREBBP* | NM_004380:c.C2399T (p.P800L) | Missense | AD | Rubinstein-Taybi syndrome 1 (180849) |  | Pat |

**Continued Supplementary Table S3. Variants of unknown significance in SNVs/InDels detected by GS**

| **Case** | **NT (mm)** | **Other sonographic findings** | **Gene** | **Mutation (all heterozygous)** | **Consequence** | **Inheritance mode** | **Disease association(s) [MIM #]** | **Pregnancy outcome** | **Inheritance confirmation** |
| --- | --- | --- | --- | --- | --- | --- | --- | --- | --- |
| 15C0520 | 6.27 | pericardial effusion | *MKKS* | NM_170784:c.G459C (p.L153F) | Missense | AR | Bardet-Biedl syndrome 6 (605231) | Live birth | Mat |
|  |  |  |  | NM_170784: c.C170T (p.T57I) | Missense | AR | Bardet-Biedl syndrome 6 (605231) | Live birth | Mat |
| 18C0028 | 4.4 | micrognathia | *PIEZO2* | NM_022068:c.A6946G (p.T2316A) | Missense | AD | ?Marden-Walker syndrome (248700) | - | Pat |
| 18C0096 | 5.3 | - | *LZTR1* | NM_006767:c.C1396T (p.R466W) | Missense | AD | Noonan syndrome 10 (616564) | - | Mat |
| 18NT0003 | 12 | Hydrosarc; cystic hygroma; short long bones | *IFT172* | NM_015662:c.A3089G(p.D1030G) | Missense | AR | Retinitis pigmentosa 71(616394); Short-rib thoracic dysplasia 10 with or without polydactyly(615630) | TOP | Pat |
|  |  |  | *IFT172* | NM_015662:c.G449C(p.G150A) | Missense | AR |  | TOP | Mat |
| 18NT0008 | 5.3 | - | *EFTUD2* | NM_004247: c.T1300A(p.C434S) | Missense | AD | Mandibulofacial dysostosis, Guion-Almeida type (610536) | Live birth | - |
| 18NT0014 | 6.6 | hydrops; cystic hygroma | *PDYN* | NM_024411:c.G638A (p.R213H) | Missense | AD | Spinocerebellar ataxia 23 (610245) | - | - |

**Continued Supplementary Table S3. Variants of unknown significance in SNVs/InDels detected by GS**

| **Case** | **NT (mm)** | **Other sonographic findings** | **Gene** | **Mutation (all heterozygous)** | **Consequence** | **Inheritance mode** | **Disease association(s) [MIM #]** | **Pregnancy outcome** | **Inheritance confirmation** |
| --- | --- | --- | --- | --- | --- | --- | --- | --- | --- |
| 18NT019 | 3.5 | - | *AHDC1* | NM_001029882:c.C1417T (p.R473C) | Missense | AD | Xia-Gibbs syndrome (615829) | - | - |
| 18NT027 | 3.5 | - | *COL2A1* | NM_001844: c.C3296T(p.P1099L) | Missense | AD | Achondrogenesis, type II or hypochondrogenesis (200610) | - | - |
|  |  |  | *LZTR1* | NM_006767:c.T2387C (p.I796T) | Missense | AD | Noonan syndrome 10 (616564) | - | - |
| 18NT028 | 3.6 | absent nasal bone | *JAG1* | NM_000214:c.C3286T(p.R1096W) | Missense | AD | Tetralogy of Fallot (187500) | Live birth | - |
| 18NT030 | 3.5 | - | *SETBP1* | NM_015559:c.C1387T(p.R463C) | Missense | AD | Mental retardation, autosomal dominant 29 (616078); Schinzel-Giedion midface retraction syndrome (269150) | - | - |
| 18C0528 | 3.5 | - | *FGFR1* | NM_023110:c.T20G(p.L7R) | Missense | AD | Hartsfield syndrome (615465) | Live birth | Mat |
| 18BA1384 | 4.0 | - | *MYH7* | NM_000257:c.C5279T(p.T1760M) | Missense | AD | Cardiomyopathy, dilated, 1S (613426) | - | - |

**Continued Supplementary Table S3. Variants of unknown significance in SNVs/InDels detected by GS**

| **Case** | **NT (mm)** | **Other sonographic findings** | **Gene** | **Mutation (all heterozygous)** | **Consequence** | **Inheritance mode** | **Disease association(s) [MIM #]** | **Pregnancy outcome** | **Inheritance confirmation** |
| --- | --- | --- | --- | --- | --- | --- | --- | --- | --- |
| 18C0588 | 3.7-4.1 | bilateral multicystic dysplastic kidneys; reverse a-wave ductus venosus; ventriculomegaly (10.5mm) | *CDON* | NM_016952:c.C2654A(p.S885X) | Stopgain | AD | Holoprosencephaly 11 (614226) | Live birth | Pat |
| 18C0905 | 6.2 | shorted cervical length (cervical incompetence) | *COL27A1* | NM_032888:c.C5462T(p.T1821I) | Missense | AR | Steel syndrome (615155) | Live birth | Mat |
|  |  |  |  | NM_032888:c.C2159T(p.P720L) | Missense | AR | Steel syndrome (615155) | Live birth | Mat |

**Supplementary Table S4. Inversions and insertions detected by GS**

| Case ID | NT (mm) | Other sonographic finding(s) or clinical indication(s) | SV(s) detected by GS | Disrupted gene(s) |
| --- | --- | --- | --- | --- |
| 17NT0004 | 4.2 | - | seq[hg19] ins(3;3)(q13.31;q21.1) g[chr3:122510925_122512050inschr3:113558845_113574795] chr3:g.113558845_113574795del | - |
| 18NT0003 | 12 | hydrosarca and short limbs | seq[hg19] ins(2;12)(q33.2;q24.31) g.[chr2:203384219_203384293inschr12:122757221_122907271cx) chr12:g.122757221_122907271del | *CLIP1*;*BMPR2* |
| 17BA0551 | 5.0 | NIPT reported a partial duplication in chromosome 8 | seq[hg19] inv(18)(q12.3) chr18:g.[38639899_38739267] | - |
| 15C0337 | 4.18 | positive Down screening risk:T21:1:3, T13 & T18 positive | seq[hg19] inv(3)(p12.2) chr3:g.[79866319_79902705] | - |
